# α-Synuclein strain homogeneity in multiple system atrophy clinical subtypes

**DOI:** 10.64898/2026.06.05.730469

**Authors:** Heather H.C. Lau, Nicholas R.G. Silver, Surabhi Mehra, Raphaella W.L. So, Le yao Li, Alison Mao, Erica Stuart, Cian Schmitt-Ulms, Bradley T. Hyman, Martin Ingelsson, Joel C. Watts

## Abstract

Conformationally distinct ‘strains’ of α-synuclein aggregates are believed to contribute to the clinical and pathological diversity observed among synucleinopathies such as multiple system atrophy (MSA) and Parkinson’s disease. Cases of MSA can be classified into two distinct clinical subtypes: the cerebellar variant, MSA-C, and the parkinsonian variant, MSA-P. To assess whether distinct α-synuclein strains may be present in individuals with MSA-C versus MSA-P, we characterized the conformational and seeding properties of α-synuclein aggregates in various brain regions from MSA-C and MSA-P patients and performed propagation studies in M83 transgenic mice. Biochemical fingerprinting of α-synuclein aggregates using limited proteolysis and a conformational stability assay failed to reveal differences between MSA-C and MSA-P either before or after propagation in mice. Similarly, using brain extracts from either MSA patients or MSA-inoculated mice, MSA-C and MSA-P α-synuclein aggregates exhibited indistinguishable seeding attributes in a seed amplification assay. Finally, no differences were observed in either the kinetics of disease progression or the extent of cerebral α-synuclein deposition in M83 mice inoculated with either MSA-C or MSA-P, regardless of the brain region from which the injected α-synuclein aggregates were derived. These results suggest that MSA clinical subtypes are unlikely to arise due to distinct α-synuclein strains. Instead, our findings support a model in which the same α-synuclein strain initially forms in different brain regions, leading to differences in disease manifestation.

## Introduction

The synucleinopathies are a group of progressive neurodegenerative disorders unified by the presence of proteinaceous inclusions within the brain that are composed of misfolded and aggregated α-synuclein (α-syn) ^1^. Examples of synucleinopathies include Parkinson’s disease (PD), dementia with Lewy bodies (DLB), and multiple system atrophy (MSA). MSA is rarer than PD but usually more aggressive, with disease progression occurring over a period of 5-6 years and death typically occurring within ten years of symptom onset ^2^. Two distinct clinical variants of MSA are recognized: the cerebellar subtype (MSA-C) and the parkinsonian subtype (MSA-P). While failure of the autonomic nervous system is common to both subtypes, MSA-P is typified by predominant parkinsonian features such as bradykinesia and rigidity that respond poorly to levodopa treatment and is associated with degeneration of the striatonigral system within the brain ^3–5^. In contrast, MSA-C presents with predominant cerebellar ataxia and gait abnormalities and is associated with degeneration of the olivopontocerebellar system. The two subtypes display regional prevalence differences, with MSA-C being more common in Japan whereas MSA-P predominates in Europe and the United States ^6–8^. MSA-P has been found to be associated with more severe cognitive dysfunction as well as a worse survival prognosis relative to MSA-C ^7,9^.

In MSA, the predominant neuropathological finding is the presence of glial cytoplasmic inclusions (GCIs) within oligodendrocytes ^10,11^, and the frequency of GCIs correlates with the severity of neurodegeneration within specific brain regions in both MSA-P and MSA-C ^5,12^. GCIs are composed of α-syn aggregates that are phosphorylated at serine-129 ^13,14^. α-Syn aggregates are self-propagating or “prion-like” because they can act as seeds that recruit monomeric α-syn into a growing fibrillar assembly ^15^. Accordingly, injection of susceptible transgenic mice or non-human primates with GCI-containing brain extracts from MSA patients induces both cerebral α-syn deposition and neurodegeneration ^16,17^. Seeding assays in cultured cells and mice have demonstrated that α-syn aggregates from MSA patient brains are more potent inducers of α-syn aggregation than samples from PD/DLB brains ^16,18–26^. One potential explanation for the variability in α-syn seeding activities between MSA and PD/DLB is the conformational strain hypothesis, which posits that structural differences between α-syn aggregates underlie the unique clinicopathologic presentations of individual synucleinopathies^27–29^. Indeed, the structure of α-syn fibrils isolated from the brains of MSA patients is distinct from the “Lewy fold” aggregate structure adopted by α-syn in the brains of individuals with PD or DLB^30,31^. Moreover, recombinant α-syn can be polymerized into structurally distinct α-syn strains that induce divergent synucleinopathies upon propagation in rodent models ^32–35^.

While the distinct clinical presentations of MSA-C and MSA-P likely stem from GCI accumulation and neurodegeneration within MSA subtype-specific brain regions, the molecular basis for how this selective vulnerability arises remains unknown. One possibility is that MSA-C and MSA-P are caused by distinct α-syn strains that preferentially target different brain regions. In the human prion disorders, phenotypic diversity and disease subtypes are known to manifest due to the presence of distinct prion strains in the brain ^36,37^, and there is evidence that clinical variants of Alzheimer’s disease may be associated with structurally distinct Aβ and/or tau aggregates ^38–43^. In support of the conformational strain theory, two different types of α-syn filaments have been found in MSA brains, although neither fibril type appears to be exclusively associated with either MSA-C or MSA-P ^30^. Moreover, it has been reported that MSA-C and MSA-P can be discriminated via a seed amplification assay (SAA) that uses α-syn aggregates from olfactory mucosa ^44^, and MSA-C- and MSA-P brains display different patterns of cellular iron deposition ^45^. Here, we tested the hypothesis that conformationally distinct strains of α-syn aggregates are present in the brains of MSA-C and MSA-P patients. Using tissue from three different brain regions, we found no evidence for MSA subtype-specific α-syn conformational variants, either before or after propagation in a transgenic synucleinopathy mouse model. Our results suggest that MSA-C and MSA-P are caused by the same α-syn strain that can initially form in different brain regions, leading to MSA subtype-specific clinical and pathological presentations.

## Results

### Conformational characterization of α-syn aggregates from MSA-C and MSA-P brains

Frozen brain tissue was obtained from three different brain regions (substantia nigra, cerebellum, and temporal cortex) from cases of MSA-C and MSA-P (**Supplemental Table S1**). The cerebellum was chosen as a potentially early site of α-syn aggregate deposition within GCIs in MSA-C whereas the substantia nigra was chosen as a potential early site of α-syn deposition within GCIs in MSA-P. The temporal cortex was selected for comparison purposes, as GCIs are infrequently observed here, whereas neuronal α-syn inclusions can be found in this region ^46^.

Initially, two cases each of MSA-C (MSA-C_1_ and MSA-C_2_) and MSA-P (MSA-P_1_ and MSA-P_2_) were obtained, which were used for the propagation studies in mice. Subsequently, six additional cases of MSA-C (MSA-C_3_ to MSA-C_8_) and five additional cases of MSA-P (MSA-P_3_ to MSA-P_7_) were obtained to supplement the conformational characterization and SAA experiments.

To characterize the conformational properties of α-syn aggregates in brain homogenates from MSA-C and MSA-P, we first utilized a conformational stability assay (CSA) that measures the relative resistance of α-syn aggregates to denaturation with guanidine hydrochloride (GdnHCl)^47^. Using the CSA, we have previously shown that MSA-associated α-syn aggregates are less stable than those present in PD or DLB brains ^20,34^. Following treatment of brain homogenates with increasing concentrations of GdnHCl, residual levels of detergent-insoluble serine-129-phosphorylated α-syn (PSyn) were assessed (**Fig. 1a**). For all three brain regions, the denaturation curves for PSyn aggregates present in the brains of MSA-C and MSA-P patients were indistinguishable (**Fig. 1b-d**). Moreover, after pooling the data from the three brain regions, the conformational stability of PSyn aggregates in MSA-C and MSA-P brains, as determined by the concentration of GdnHCl required to solubilize 50% of the aggregates (the [GdnHCl]_50_ value), were essentially identical (**Fig. 1e**). We also performed conformational fingerprinting of α-syn aggregates in brain homogenates from MSA-C and MSA-P patients using protease digestion. α-Syn aggregates are partially resistant to protease digestion and, due to structural differences, distinct α-syn aggregate strains can produce unique banding patterns following digestion with proteases such as thermolysin (TL) or proteinase K (PK) ^34,35,48^. While some case-to-case variability was present in the patterns of TL- and PK-resistant α-syn species, no consistent differences were observed between samples from MSA-C versus MSA-P patients for any of the three brain regions (**Fig. 2**).

**Figure 1.**
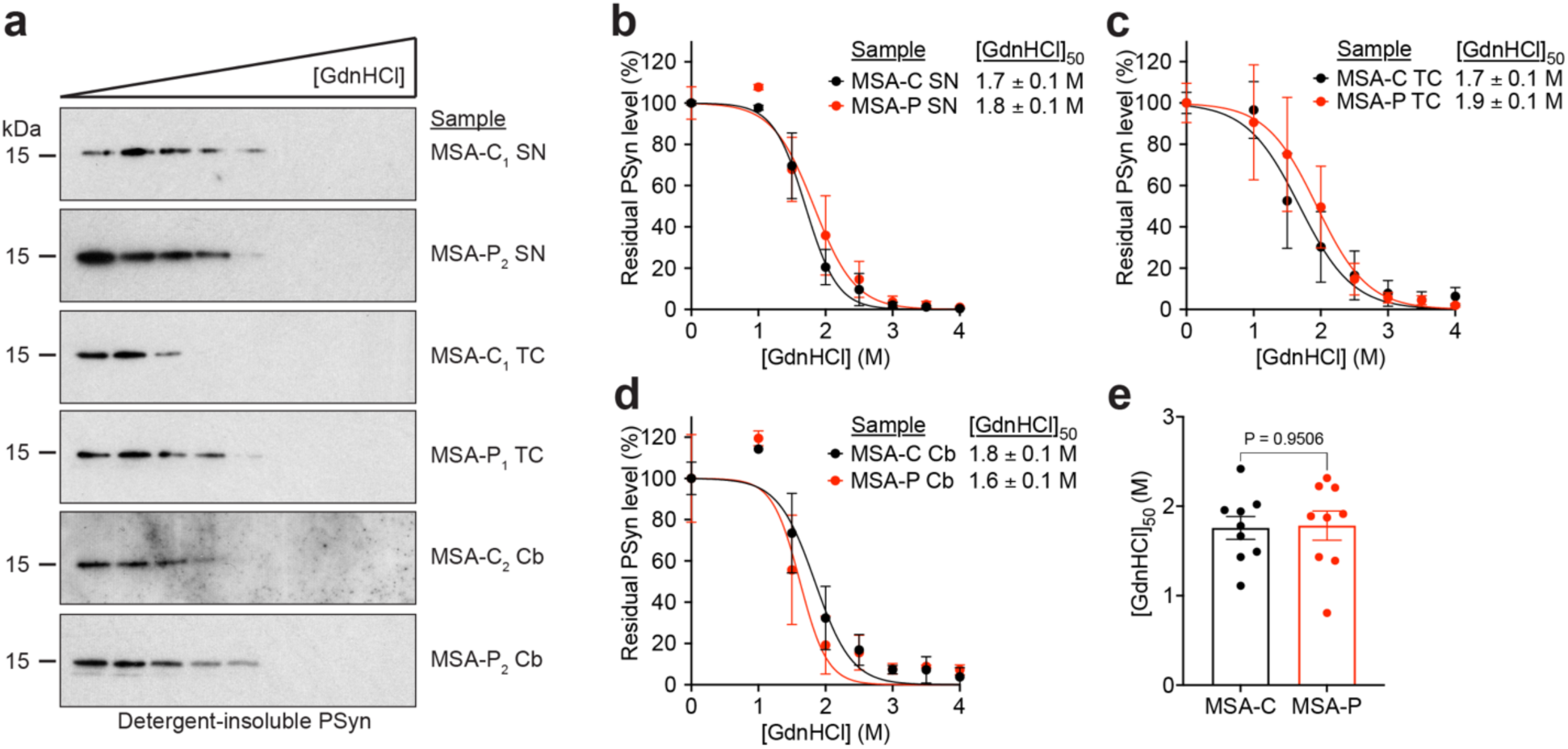
Conformational stability of α-syn aggregates from MSA-C and MSA-P patient brain samples. **a**) Representative immunoblots of residual levels of detergent-insoluble PSyn species following treatment of samples from different brain regions of MSA-C and MSA-P patients with increasing concentrations of GdnHCl. PSyn was detected using the antibody EP1536Y. SN: substantia nigra; TC: temporal cortex; Cb: cerebellum. **b**-**d**) GdnHCl denaturation curves for PSyn aggregates in samples from the substantia nigra (b), temporal cortex (c), and cerebellum (d) of MSA-C and MSA-P patients. Data is mean ± s.e.m. for n = 3 independent samples per brain region and MSA subtype. **e**) Combined [GdnHCl]_50_ values for PSyn aggregates in MSA-C or MSA-P patient brain samples (data is mean ± s.e.m.; n = 9 for each of MSA-C and MSA-P). Statistical significance was assessed using a Mann-Whitney test.

**Figure 2.**
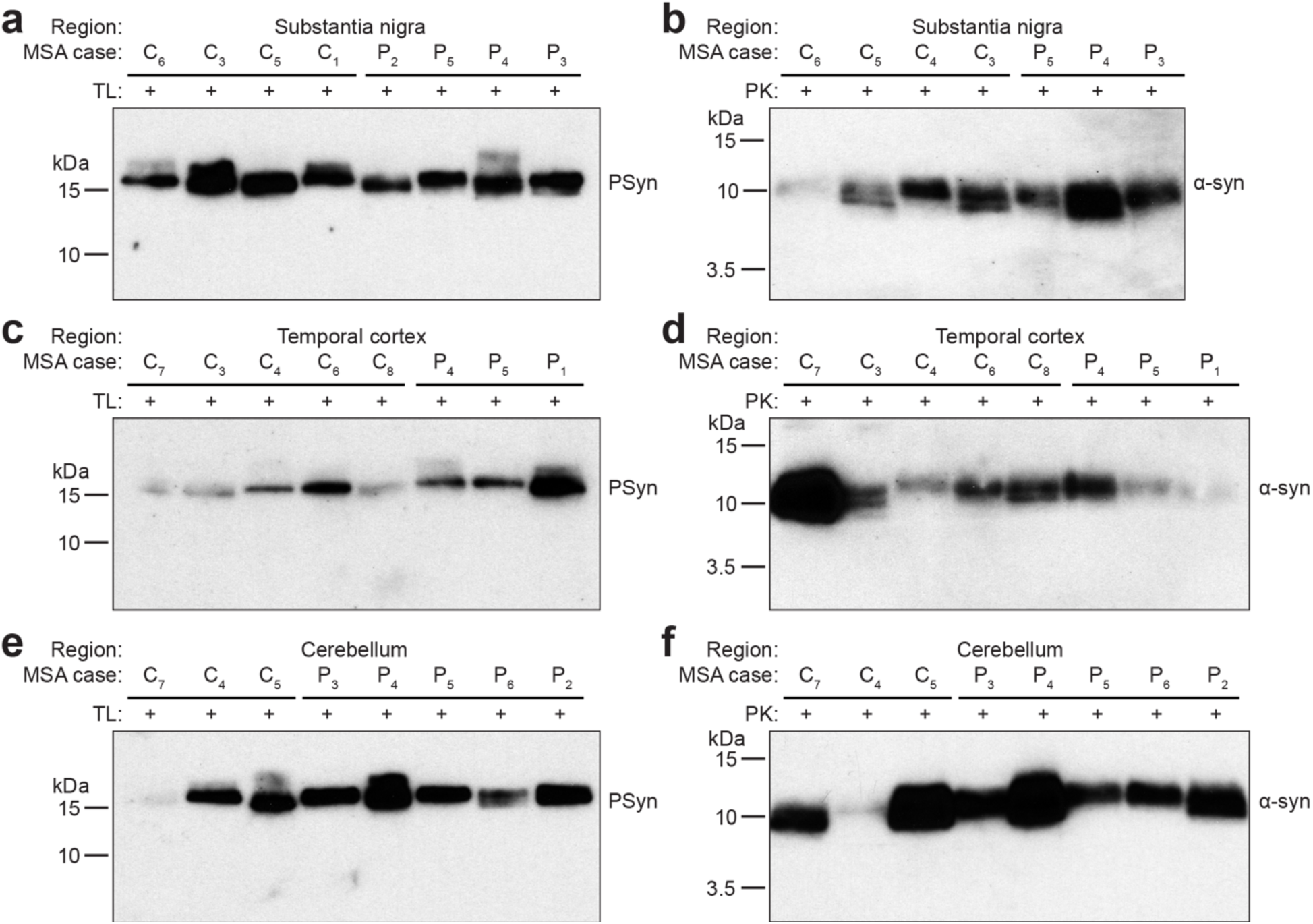
Conformational fingerprinting of α-syn aggregates from MSA-C and MSA-P patient brain samples using protease digestion. Immunoblots of detergent-insoluble PSyn or α-syn species following digestion of samples from either the substantia nigra (a, b), temporal cortex (c, d), or cerebellum (e, f) of MSA-C and MSA-P patients with thermolysin (TL; a, c, e) or proteinase K (PK; b, d, f). PSyn was detected using the antibody EP1536Y and α-syn was detected using the antibody Syn-1.

### Propagation of α-syn aggregates from MSA-C and MSA-P brains in M83^+/-^ mice

Intracerebral injection of M83 transgenic mice, which express A53T-mutant human α-syn, with brain homogenate from MSA patients causes neurological illness, as indicated by the presence of progressive motor impairment accompanied by a cerebral synucleinopathy ^16,18,56^. Thus, we decided to compare the propagation properties of α-syn aggregates from MSA-C and MSA-P patients upon inoculation into M83 mice. For these experiments, hemizygous M83 mice (M83^+/-^) were used because they do not develop spontaneous disease prior to 22 months of age ^56^.

M83^+/-^ mice at ∼5 weeks of age were intracerebrally inoculated with homogenate of different brain regions from two cases each of MSA-C and MSA-P, and then monitored for the development of motor impairment over a period of 400-500 days post-inoculation (**Fig. 4a**, **Table 1**). Considerable variability in the amount of TL-resistant, detergent-insoluble PSyn aggregates was present in the samples used for inoculation (**Fig. 4b**). However, all samples transmitted disease to M83^+/-^ mice, albeit with varying efficiency (**Table 1**). For samples derived from the substantia nigra or cerebellum, 32/33 and 15/16 injected mice respectively developed motor impairment. Transmission efficiency was lower when using temporal cortex samples, with only 23/33 mice exhibiting signs of neurological illness, likely due to a lower quantity of GCIs in this brain region. While samples from MSA-C patients tended to produce disease in M83^+/-^ mice slightly more rapidly than samples from MSA-P patients (**Fig. 4c-e**), no significant difference in the survival curves were observed for MSA-C-inoculated mice versus MSA-P-inoculated mice when the data for all brain regions was pooled (**Fig. 4f**). Following inoculation with MSA homogenate, the disease incubation times (i.e., the time from inoculation to euthanasia due to progressive motor impairment) were not significantly different between male and female M83^+/-^mice (**Fig. 4g**).

**Table 1.**
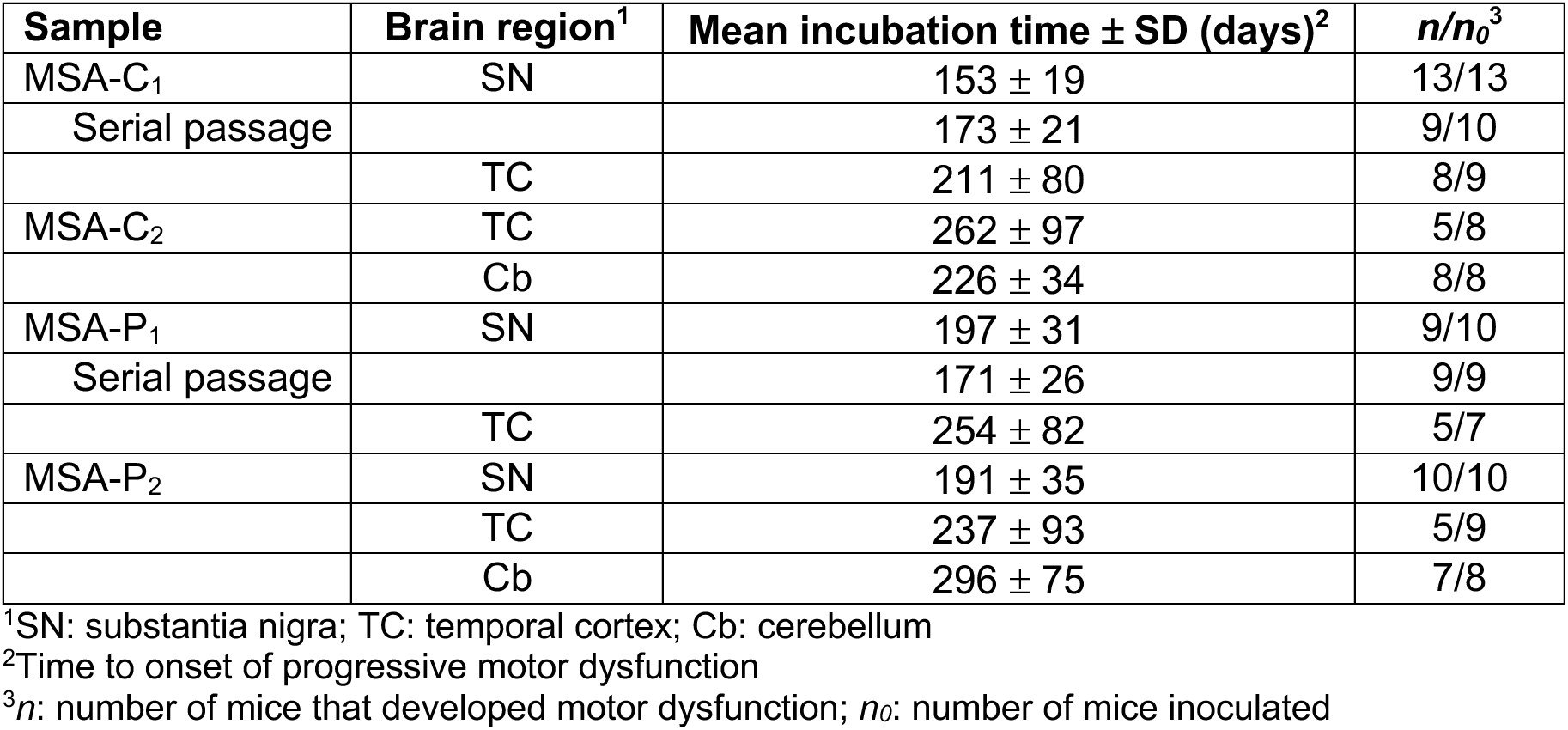
Incubation times for M83^+/-^ mice inoculated with MSA-C or MSA-P brain extract.

| Sample | Brain region<sup>1</sup> | Mean incubation time ± SD (days)<sup>2</sup> | $n/n_0$<sup>3</sup> |
| --- | --- | --- | --- |
| MSA-C<sub>1</sub> | SN | 153 ± 19 | 13/13 |
| Serial passage | | 173 ± 21 | 9/10 |
| | TC | 211 ± 80 | 8/9 |
| MSA-C<sub>2</sub> | TC | 262 ± 97 | 5/8 |
| | Cb | 226 ± 34 | 8/8 |
| MSA-P<sub>1</sub> | SN | 197 ± 31 | 9/10 |
| Serial passage | | 171 ± 26 | 9/9 |
| | TC | 254 ± 82 | 5/7 |
| MSA-P<sub>2</sub> | SN | 191 ± 35 | 10/10 |
| | TC | 237 ± 93 | 5/9 |
| | Cb | 296 ± 75 | 7/8 |

<sup>1</sup>SN: substantia nigra; TC: temporal cortex; Cb: cerebellum
<sup>2</sup>Time to onset of progressive motor dysfunction
<sup>3</sup> $n$ : number of mice that developed motor dysfunction; $n_0$ : number of mice inoculated

Next, we wondered whether α-syn aggregates in brain homogenates from MSA-C and MSA-P cases can be distinguished using α-syn SAA. In this assay, α-syn aggregate seeds present in cerebrospinal fluid or other biological samples stimulate the aggregation of recombinant α-syn, which can be detected in real-time in a microplate reader using the aggregate-binding fluorescent dye thioflavin T (ThT) ^49^. SAAs have previously been used to discriminate between PD and MSA samples, based on the maximum ThT fluorescence signal produced ^50–52^. Whether α-syn SAA can distinguish MSA-C and MSA-P remains unresolved and may depend on the biological source of α-syn seeds ^44,53^. α-Syn SAAs were performed using recombinant wild-type human α-syn purified using an osmotic shock protocol, which greatly reduces spontaneous aggregation of α-syn and improves SAA performance ^54^, as well as a reaction buffer optimized for amplification of MSA-associated α-syn aggregates ^52^. As has been previously reported ^52^, there was significant heterogeneity in the observed seeding behavior for individual MSA samples (**Supplemental Fig. S1**). However, when the seeding data for all cases of a given MSA subtype were pooled and analyzed collectively, the amplification curves were essentially superimposable, and no statistically significant MSA subtype-specific differences were found for any of the SAA parameters examined (lag phase, maximum ThT fluorescence signal, or plateau ThT fluorescence signal) (**Fig. 3**). The same result was obtained regardless of whether homogenate from the substantia nigra, temporal cortex, or cerebellum of MSA-C and MSA-P cases was used as the seed. On average, the ThT fluorescence signals were lower when using temporal cortex samples, likely because the quantity of GCIs is lower in this brain region ^55^. Collectively, we were unable to find any biochemical or *in vitro* seeding differences between α-syn aggregates from MSA-C and MSA-P patients.

**Figure 3.**
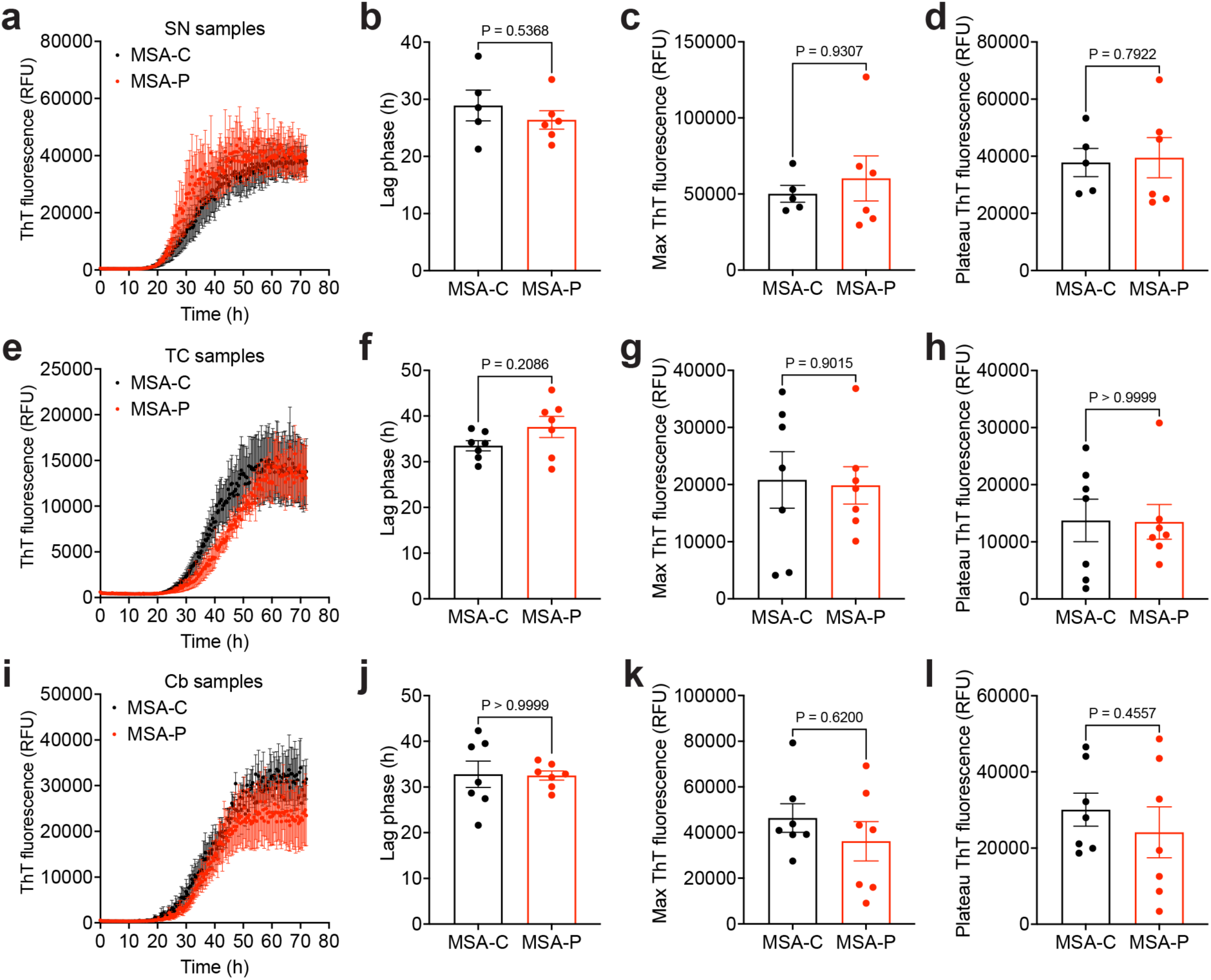
α-Synuclein seed amplification assays on MSA-C and MSA-P patient brain samples. **a, e, i**) ThT aggregation curves for SAAs using homogenates from the substantia nigra (a), temporal cortex (e), or cerebellum (i) of MSA-C and MSA-P patients. **b, f, j**) Calculated lag phases for SAAs utilizing substantia nigra (b), temporal cortex (f), or cerebellum (j) tissue. **c, g, k**) Maximum ThT fluorescence values for SAAs utilizing substantia nigra (c), temporal cortex (g), or cerebellum (k) tissue. **d, h, l**) Plateau ThT fluorescence values for SAAs utilizing substantia nigra (d), temporal cortex (h), or cerebellum (l) tissue. For SN samples, n = 5 (MSA-C) or n = 6 (MSA-P); for TC and Cb samples, n = 7 per MSA subtype. For each sample, 3-4 technical replicates were performed. Data is mean ± s.e.m. Statistical significance was assessed using a Mann-Whitney test.

**Figure 4.**
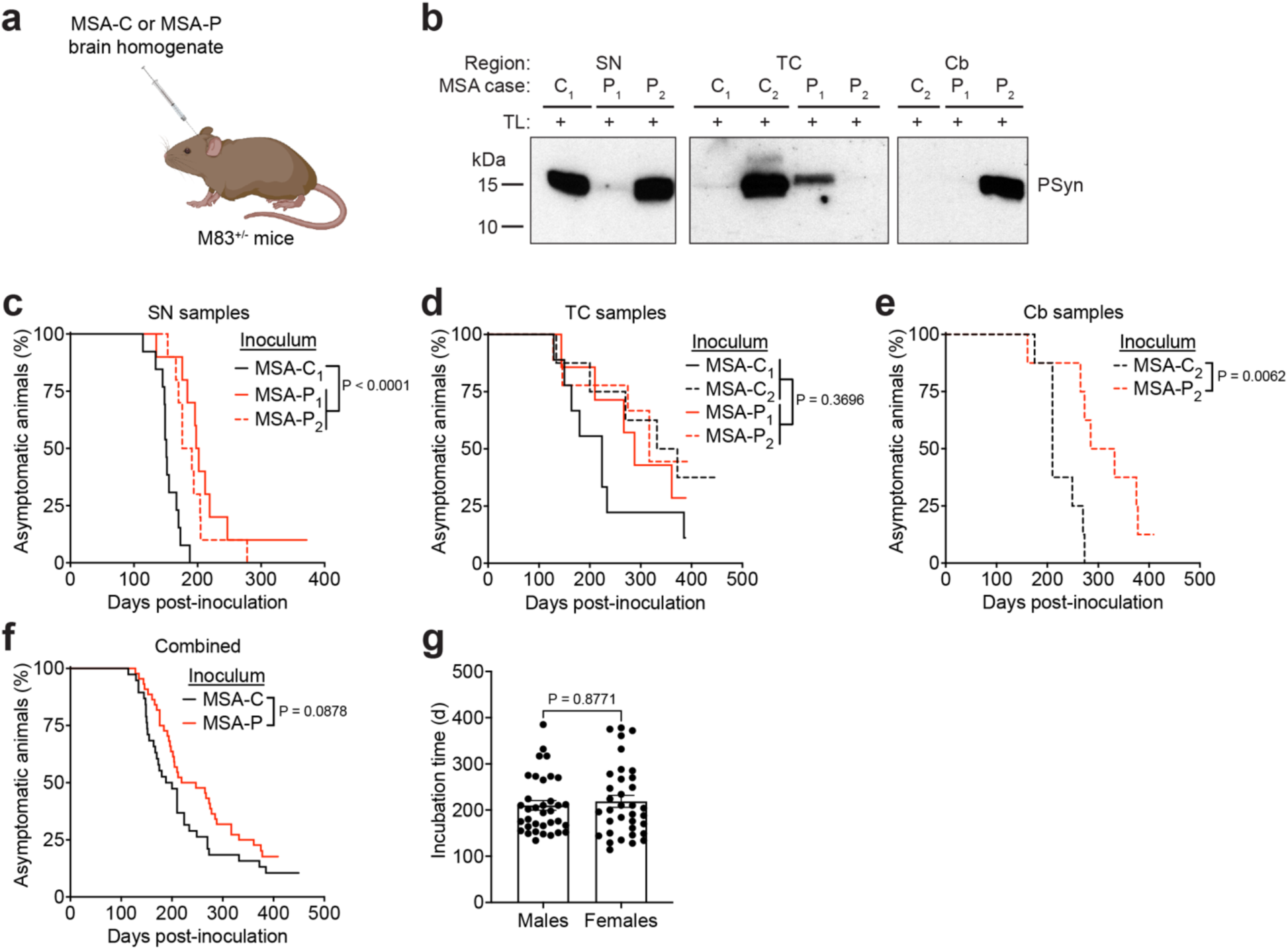
Propagation of MSA-C and MSA-P α-syn aggregates in M83^+/-^ mice. **a**) Schematic of the intracerebral inoculation experiments in M83^+/-^ mice. **b**) Immunoblots showing relative amounts of thermolysin-resistant, detergent-insoluble PSyn species in the MSA samples used for inoculation. PSyn was detected using the antibody EP1536Y. **c**-**e**) Kaplan-Meier survival curves for M83^+/-^ mice inoculated with homogenates of the substantia nigra (c), temporal cortex (d), or cerebellum (e) from MSA-C or MSA-P patients. For SN samples, n = 13, 10, or 10 mice for MSA-C_1_, MSA-P_1_, and MSA-P_2_, respectively; for TC samples, n = 9, 8, 7, or 9 mice for MSA-C_1_, MSA-C_2_, MSA-P_1_, and MSA-P_2_, respectively; for Cb samples, n = 8 for both MSA-C_2_ and MSA-P_2_. **f**) Combined Kaplan-Meier survival curve for all M83^+/-^ mice inoculated with samples from MSA-C or MSA-P patients (n = 38 for mice inoculated with MSA-C, n = 44 for mice inoculated with MSA-P). In panels c-f, statistical significance was assessed using the Log-rank test. **g**) Incubation times for MSA-inoculated male and female M83^+/-^ mice. Data is mean ± s.e.m. for n = 35 per sex. Only mice that developed motor dysfunction were included. Statistical significance was assessed using a Mann-Whitney test.

To conformationally characterize the induced α-syn aggregates in the brains of MSA-inoculated M83^+/-^ mice, we first performed CSAs. As with the human MSA samples, no consistent differences in α-syn aggregate stability were observed between MSA-C-inoculated mice and MSA-P-inoculated mice, regardless of which brain region was used as the inoculum (**Fig. 5a-e**). Similarly, the banding patterns of PK- and TL-resistant α-syn aggregates in the brains of M83^+/-^mice inoculated with either MSA-C or MSA-P were indistinguishable (**Fig. 5f**). Finally, using an identical protocol as used for the human MSA samples, we performed α-syn SAAs on brain homogenates from M83^+/-^ mice inoculated with substantia nigra homogenate from MSA-C or MSA-P patients. As expected, samples from clinically ill MSA-C-inoculated mice and MSA-P-inoculated mice were positive by SAA, whereas samples from asymptomatic buffer-inoculated M83^+/-^ mice did not produce an increase in ThT fluorescence (**Fig. 6a**). No statistically significant differences in either the lag phase, maximum ThT fluorescence, or plateau ThT fluorescence values were observed for samples from mice inoculated with MSA-C versus mice inoculated with MSA-P (**Fig. 6b-d**). In summary, the conformational attributes of induced α-syn aggregates in the brains of M83^+/-^ mice inoculated with either MSA-C or MSA-P are highly similar, if not identical.

**Figure 5.**
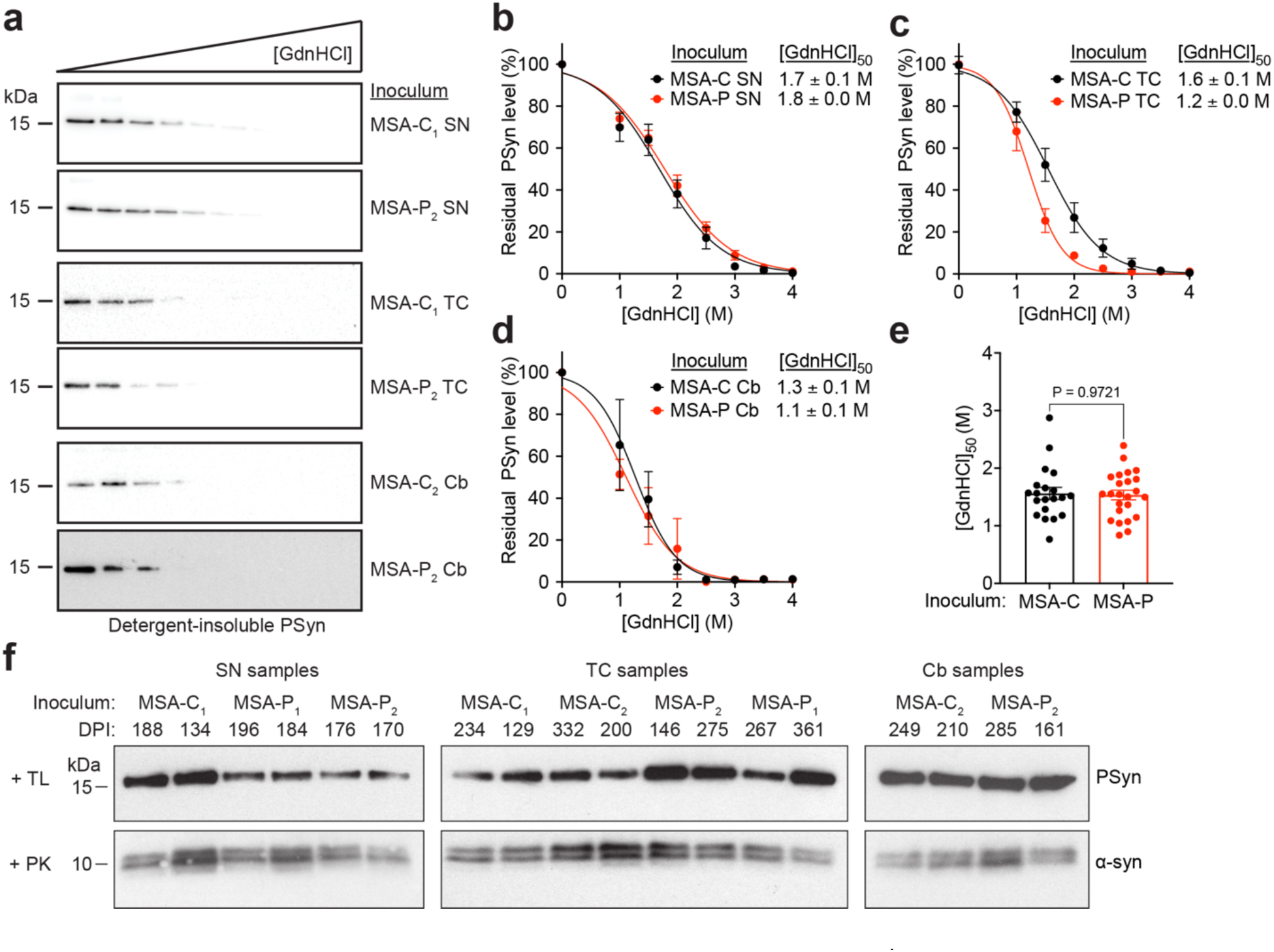
Conformational analysis of α-syn aggregates in M83^+/-^ mice inoculated with MSA-C or MSA-P brain extract. **a**) Representative immunoblots of residual levels of detergent-insoluble PSyn species following treatment of brain homogenates from clinically ill M83^+/-^ mice that were inoculated with the indicated MSA samples with increasing concentrations of GdnHCl. **b**-**d**) GdnHCl denaturation curves for PSyn aggregates in samples from the brains of M83^+/-^mice inoculated with extracts from the substantia nigra (b), temporal cortex (c), or cerebellum (d) of MSA-C and MSA-P patients. Data is mean ± s.e.m. For SN samples, n = 7 and 14 mice for MSA-C and MSA-P, respectively; for TC samples, n = 10 and 7 mice for MSA-C and MSA-P, respectively; for Cb samples, n = 3 mice for both MSA-C and MSA-P. **e**) Combined [GdnHCl]_50_ values for M83^+/-^ mice inoculated with samples derived from different brain regions of MSA-C or MSA-P patients (n = 20 for mice inoculated with MSA-C, n = 24 for mice inoculated with MSA-P). Statistical significance was assessed using a Mann-Whitney test. **f**) Immunoblots of detergent-insoluble PSyn or α-syn species following proteinase K or thermolysin digestion of brain homogenates from clinically ill M83^+/-^ mice at the indicated days post-inoculation (DPI) with the specified MSA samples. In panels a and f, PSyn was detected using the antibody EP1536Y and total α-syn was detected using the antibody Syn-1.

**Figure 6.**
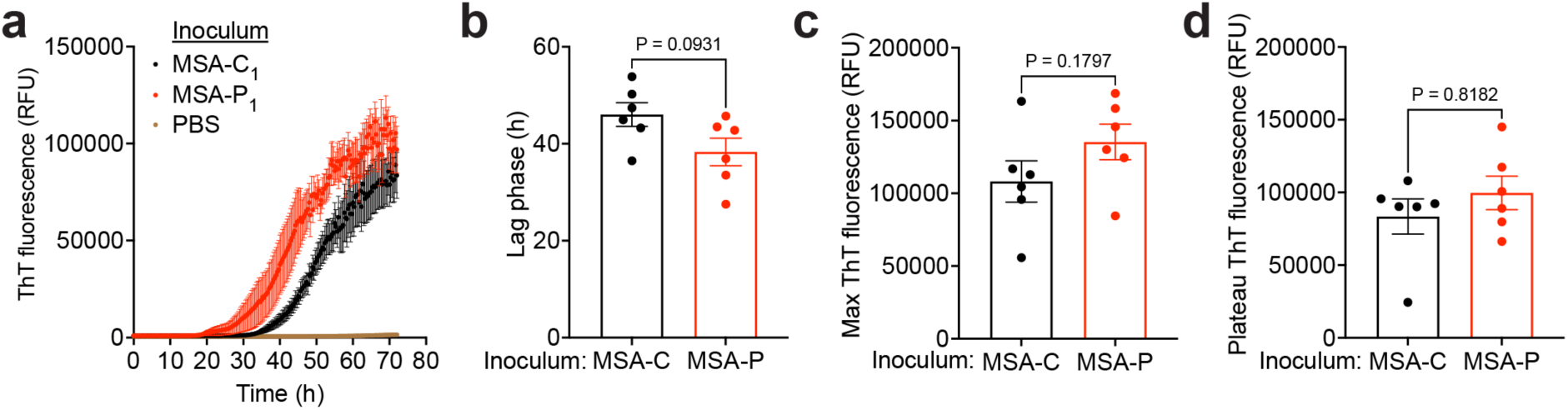
α-Synuclein seed amplification assays on brain homogenates from M83^+/-^ mice inoculated with MSA-C or MSA-P. **a**) ThT aggregation curves for SAAs using brain homogenates from clinically ill M83^+/-^ mice inoculated with substantia nigra homogenate from either MSA-C or MSA-P patients (n = 6 mice per MSA subtype). As a negative control, brain homogenates from asymptomatic M83^+/-^ mice at 5 months post-inoculation with PBS were used (n = 5). **b**) Calculated lag phases for SAAs. **c**) Maximum ThT fluorescence values for SAAs. **d**) Plateau ThT fluorescence values for SAAs. For each sample, 3-4 technical replicates were performed. Data is mean ± s.e.m. Statistical significance was assessed using a Mann-Whitney test.

We have previously shown that various recombinant and brain-derived α-syn strains can produce distinct neuropathological signatures upon propagation in M83^+/-^ mice ^20,34,35,48^. Thus, we examined the distribution pattern of PSyn deposits in the brains of M83^+/-^ mice inoculated with either MSA-C or MSA-P. As previously reported ^16,34^, the PSyn deposits in MSA-inoculated mice were exclusively present in neurons and exhibited a “ring-like” morphology, and this did not differ dependent on the brain region or MSA subtype from which the inoculum was derived (**Fig. 7a**). The induced PSyn deposits in the brains of MSA-inoculated M83^+/-^ mice were primarily located in regions of the hindbrain such as the hypothalamus, midbrain, and brainstem, with less frequent deposits in the cortex. There were no consistent differences in the regional targeting of PSyn deposits between mice inoculated with samples from different brain regions of either MSA-C or MSA-P (**Fig. 7b, c, Supplemental Fig. S2**). Thus, MSA-C and MSA-P produce identical neuropathological signatures upon propagation in M83^+/-^ mice.

**Figure 7.**
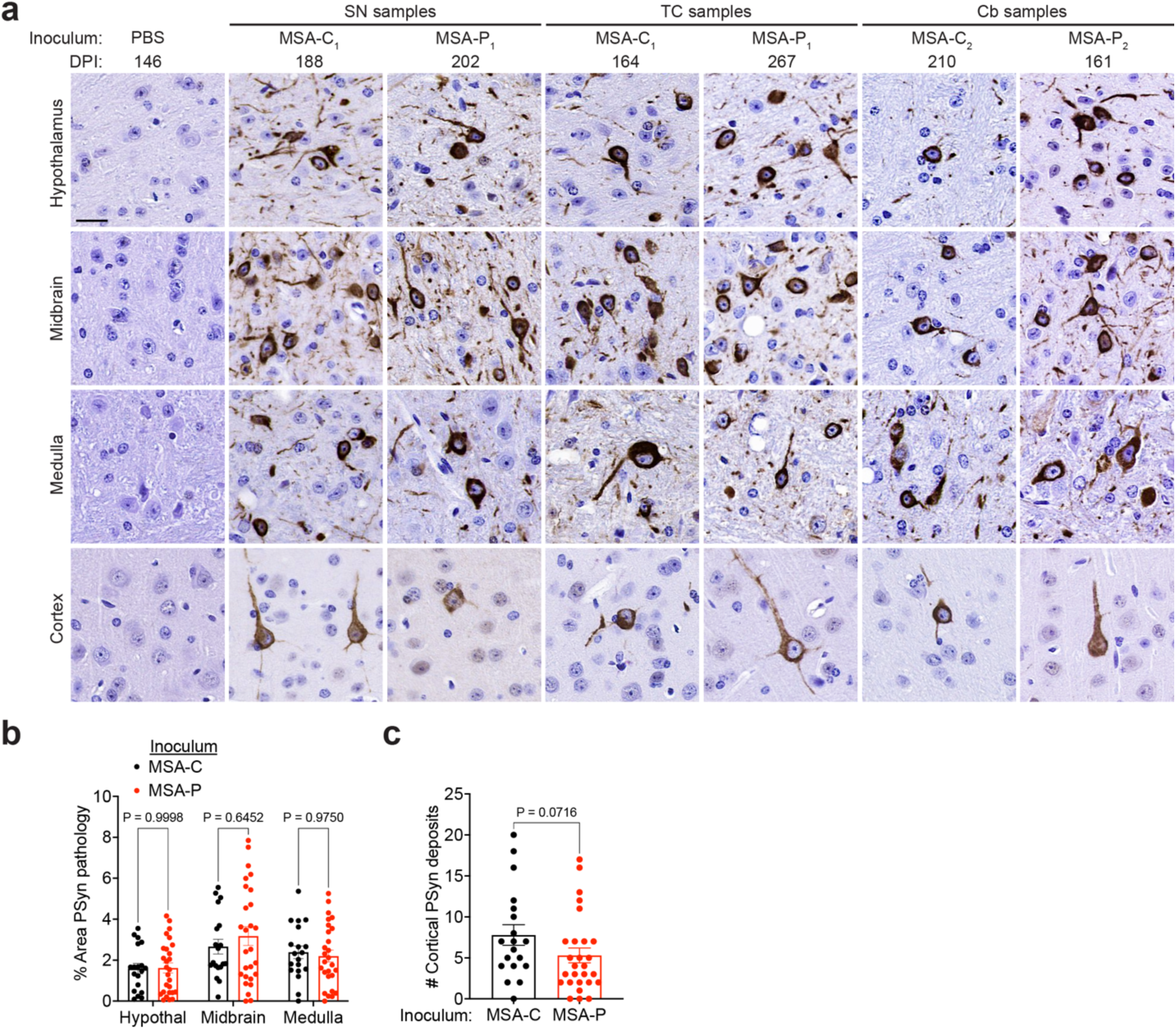
Neuropathological analysis of M83^+/-^ mice inoculated with MSA-C or MSA-P brain extract. **a**) Representative images of PSyn-stained sections from the hypothalamus, midbrain, medulla, and cortex of clinically ill M83^+/-^ mice following inoculation with the indicated MSA samples. Sections from the brain of an asymptomatic M83^+/-^ mouse following inoculation with PBS are shown as a negative control. PSyn was detected using the antibody EP1536Y. Scale bar = 20 µm (applies to all sections). **b**) Quantification of the percent area covered by PSyn staining in the indicated brain regions from MSA-inoculated M83^+/-^ mice. Statistical significance was assessed by two-way ANOVA followed by Šidák’s multiple comparisons test. **c**) Quantification of the number of cortical PSyn deposits in the brains of MSA-inoculated M83^+/-^mice. Statistical significance was assessed using a Mann-Whitney test. In panels b and c, data for SN, TC, and Cb inocula are pooled for each MSA subtype. Graphs display mean ± s.e.m for n = 19 (MSA-C) and n = 27 (MSA-P) inoculated M83^+/-^ mice.

Lastly, we asked whether subtle differences in the pathological properties of α-syn aggregates in M83^+/-^ mice inoculated with MSA-C versus MSA-P may only become apparent upon secondary (serial) passaging. Brain homogenates from clinically ill M83^+/-^ mice previously inoculated with substantia nigra homogenate from either MSA-C or MSA-P were intracerebrally inoculated into naïve M83^+/-^ mice (**Fig. 8a**). No differences in the kinetics of disease development were observed between mice inoculated with passaged MSA-C or MSA-P (**Fig. 8b**, **Table 1**). Moreover, using the CSA or by performing protease digestion assays using PK and TL, no differences in the conformational profiles of α-syn aggregates in the brains of mice inoculated with passaged MSA-C or MSA-P were observed (**Fig. 8c-f**). Finally, the distribution pattern of PSyn aggregates in the brains of mice inoculated with either of the two passaged MSA subtypes was highly similar (**Fig. 8g, h, Supplemental Fig. S3**). These results demonstrate that serial passage of α-syn aggregates from MSA-C and MSA-P patients in M83^+/-^mice produces a consistent molecular and neuropathological phenotype, with no apparent MSA subtype-specific differences.

**Figure 8.**
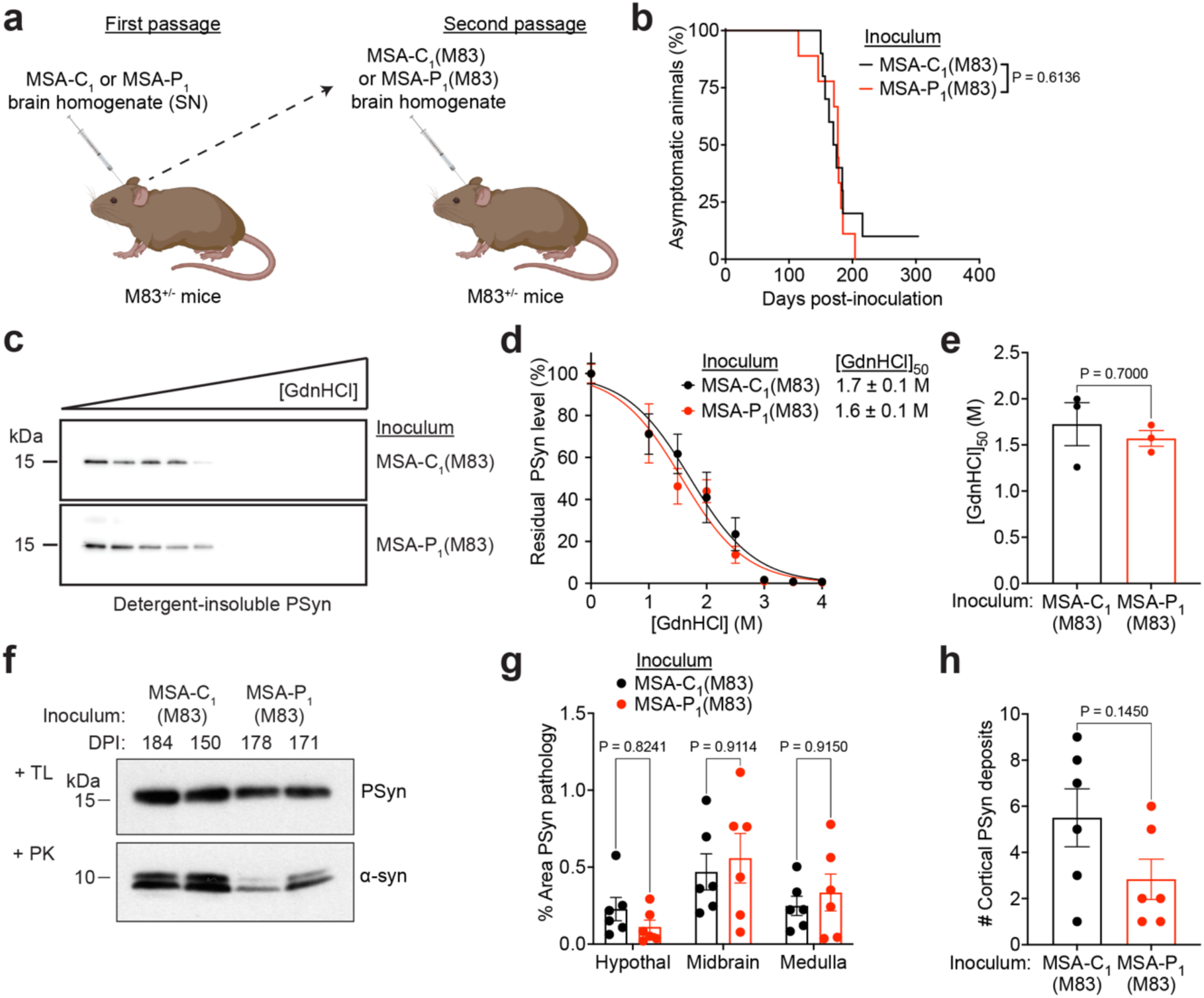
Serial passage of MSA-P and MSA-C α-syn aggregates in M83^+/-^ mice. **a**) Schematic of serial propagation experiments in M83^+/-^ mice using brain samples from animals originally injected with substantia nigra homogenate from the MSA-C_1_ [MSA-C_1_(M83)] or MSA-P_1_ [MSA-P_1_(M83)] cases. **b**) Kaplan-Meier survival curves for M83^+/-^ mice inoculated with MSA-C_1_(M83) (n = 10) or MSA-P_1_(M83) (n = 9) brain homogenate. Statistical significance was determined using the Log-rank test. **c**) Representative immunoblots of residual levels of detergent-insoluble PSyn species following treatment of brain homogenates from clinically ill M83^+/-^ mice that were inoculated with the indicated passaged MSA samples with increasing concentrations of GdnHCl. **d, e**) GdnHCl denaturation curves (d) and [GdnHCl]_50_ values (e) for PSyn aggregates in samples from the brains of M83^+/-^ mice inoculated with passaged MSA-C and MSA-P extracts. Data is mean ± s.e.m. for n = 3 mice each for MSA-C_1_(M83) and MSA-P_1_(M83). Statistical significance was assessed using a Mann-Whitney test. **f**) Immunoblots of detergent-insoluble PSyn or α-syn species following PK or TL digestion of brain homogenates from M83^+/-^ mice at the indicated days post-inoculation with the specified passaged MSA samples. PSyn was detected using the antibody EP1536Y and total α-syn was detected using the antibody Syn-1. **g**) Quantification of the percent area covered by PSyn staining in the indicated brain regions from M83^+/-^ mice inoculated with passaged MSA brain extracts. Statistical significance was assessed by two-way ANOVA followed by Šidák’s multiple comparisons test. **h**) Quantification of the number of cortical PSyn deposits in the brains of M83^+/-^ mice inoculated with passaged MSA brain extracts. Statistical significance was assessed using a Mann-Whitney test. In panels g and h, graphs display mean ± s.e.m for n = 6 mice each for MSA-C_1_(M83) and MSA-P_1_(M83).

## Discussion

Conformational strains of protein aggregates are increasingly recognized as drivers of clinical and pathological heterogeneity in human neurodegenerative diseases ^28,57–59^. Two potential scenarios seemed plausible for explaining the existence of phenotypic heterogeneity between MSA clinical subtypes. In the first scenario, distinct strains of α-syn aggregates are present in MSA-C versus MSA-P, each of which exhibits an initial tropism for specific brain regions. For instance, the strain that causes MSA-C may exhibit an initial preference for replicating in the olivopontocerebellar system, leading to GCI formation and cell loss in the cerebellum and a clinical presentation dominated by cerebellar symptoms ^60^. In contrast, the strain that causes MSA-P may preferentially replicate in brain regions such as the substantia nigra at first, leading to GCIs, nigral cell loss, and parkinsonian symptoms. Indeed, α-syn strains are known to propagate in specific brain regions and cell types in a synucleinopathy mouse model ^34^. In the second scenario, the same α-syn strain is present in both MSA-C and MSA-P. In this situation, the distinct clinical presentations may result from the initial, potentially random formation of a common strain in different brain regions (i.e., the strain may initially form either in the cerebellum or basal ganglia, respectively causing the MSA-C and MSA-P clinical subtypes).

Our data strongly suggest that MSA-C and MSA-P are caused by the same α-syn strain. No differences in the conformational stability of α-syn aggregates, the structural fingerprint of α-syn aggregates following limited proteolysis, or the seeding attributes during α-syn SAA were observed between MSA-C and MSA-P, either before or after propagation of α-syn aggregates in M83^+/-^ mice. In line with these observations, neither the kinetics of disease progression nor the resultant patterns of cerebral α-syn deposition differed between M83^+/-^ mice that were inoculated with either MSA-C or MSA-P. As different α-syn strains, derived either from recombinant α-syn polymerized into fibrils or brain extracts from synucleinopathy-laden mouse or human brains, produce biochemically and neuropathologically distinct synucleinopathies upon propagation in M83^+/-^ mice ^20,34,35,48^, our transmission studies do not support the hypothesis that MSA-C and MSA-P are caused by different α-syn strains. Moreover, samples from three different brain regions of MSA patients, including regions specifically linked to the initial appearance of GCIs in either MSA-C or MSA-P, each produced an indistinguishable synucleinopathy in M83^+/-^ mice.

Collectively, these results support a model in which the same α-syn strain initially forms stochastically in either the cerebellum or the substantia nigra, respectively leading to the MSA-C and MSA-P clinical subtypes. As the disease progresses, the α-syn aggregates spread away from their original site of formation, eventually leading to a partial convergence of clinical and pathological presentations.

Conformational strains are associated with structurally distinct protein aggregates, differing in factors such as the fold of the monomeric protein unit within a protofilament and/or the number and arrangement of protofilaments within a fibrillar aggregate ^61^. Cryo-EM studies have revealed that two distinct α-syn fibrillar structures (type I and type II) can be found in MSA patient brains, each of which consists of two asymmetric protofilaments ^30^. Subvariants of both type I and type II filaments can be discerned in MSA brains, and there is heterogeneity in the relative proportion of the various filament types within a single brain ^30,62^. Based on the limited number of cases that have been examined so far, type I or type II filaments do not seem to be preferentially associated with either MSA-C or MSA-P ^30^, which is consistent with our observation of α-syn strain homogeneity between MSA clinical subtypes. Because the structures of type I and type II MSA α-syn filaments are more closely related to each other than they are to the Lewy fold filament structure observed in PD/DLB brains ^31^, it is possible that our biochemical and seeding assays were not sufficiently granular to permit discrimination between potentially related but distinct α-syn aggregate structures in MSA-C and MSA-P. Nonetheless, even if subtle conformational variations in α-syn aggregates between MSA-C and MSA-P do exist, our seeding and transmission studies suggest that they are unlikely to confer a major functional difference. While our findings suggest that MSA-C and MSA-P are unlikely to be caused by distinct α-syn strains, this is not to say that all MSA cases are caused by a single unique strain of α-syn aggregates. Indeed, there are several examples of atypical MSA variants, both at the clinical and pathological level ^63^. While distinct α-syn fibril structures have been found in one type of atypical MSA and juvenile-onset synucleinopathy ^64,65^, the ‘typical’ MSA fibril structure was found in an MSA variant with frontotemporal lobar degeneration ^66^. Other experimental approaches are unearthing additional potential MSA pathological subtypes that could conceivably be rooted in the presence of different α-strains ^67,68^.

Although brain extracts from MSA patients reproducibly induce a cerebral synucleinopathy upon injection into M83^+/-^ transgenic mice, it must be noted that the resultant disease does not accurately mimic MSA ^16,18,69^. Most notably, while most α-syn inclusions in MSA are found in oligodendrocytes, α-syn deposits in MSA-inoculated M83^+/-^ mice are almost exclusively found in neurons. The reasons for this discrepancy remain unclear. It is unlikely to be related to the expression of A53T-mutant human α-syn in M83^+/-^ mice since transgenic mice expressing wild-type human α-syn also develop predominantly neuronal α-syn inclusions following inoculation with MSA brain extract containing α-syn aggregates composed of sequence-matched wild-type human α-syn ^70^. While the mouse prion protein promoter used to drive expression of the A53T-mutant human α-syn transgene in M83^+/-^ mice does favor gene expression in neurons and astrocytes over oligodendrocytes ^71^, α-syn expression within improper cell types or brain regions in M83^+/-^ mice is also unlikely to explain the discrepancy since transgenic mice expressing A53T-mutant human α-syn driven by the α-syn promoter develop mostly neuronal and astroglial α-syn inclusions following MSA inoculation ^72^. Mouse oligodendrocytes can support α-syn aggregation as transgenic mice expressing α-syn driven by an oligodendrocyte-specific promoter develop α-syn inclusions following injection with recombinant α-syn strains ^73^. One possibility is that α-syn inclusions in MSA and MSA-inoculated M83^+/-^ mice initially develop in neurons and then form later in the disease process in oligodendrocytes, possibly via direct transfer of aggregate seeds from neighboring neurons ^74^. In M83^+/-^ mice, potentially due to high levels of α-syn overexpression in spinal cord neurons ^56^, accumulation of neuronal α-syn aggregates may elicit severe motor dysfunction before α-syn aggregates have had a chance to spread to oligodendrocytes. New evidence does indeed suggest that neuronal α-syn inclusions are a major driver of neurodegeneration in MSA ^75^, and mice injected with recombinant α-syn fibrils or an MSA-like recombinant α-syn strain do exhibit α-syn inclusions in oligodendrocytes later in the disease course ^76,77^.

Due to the prion-like nature of α-syn aggregates, it is generally assumed that α-syn aggregate structures are faithfully propagated upon inoculation of mice. For example, the differential conformational stabilities of α-syn aggregates from MSA versus PD patients are maintained upon propagation in M83^+/-^ mice ^20^. However, recent data suggests that the unique asymmetric two-protofilament structure that defines α-syn aggregates from MSA patient brains is not conserved following injection of M83^+/-^ mice with MSA patient brain extract ^78^. Specifically, α-syn fibrils in MSA-inoculated M83^+/-^ mice consist of a single protofilament that most closely resembles a type II MSA protofilament subvariant, not the asymmetric two-protofilament type I fibrils found in the original MSA brain. Consistent with this finding, the structures of an MSA-like recombinant α-syn fibril strain before and after propagation in M83^+/-^ mice are similar, but not identical to each other, and neither structure accurately recapitulates the structure of brain-derived MSA α-syn aggregates ^77^. Following amplification of MSA-derived α-syn aggregates by SAA, the amplified and original MSA aggregates induce an identical disease in M83^+/-^ mice, arguing for structural maintenance during seeding ^79^. Although the SAA-amplified α-syn aggregates also consist of two protofilaments, each protofilament adopts the same molecular fold and the two protofilaments are arranged symmetrically rather than in the asymmetric arrangement observed in MSA brain-derived fibrils ^30,79^. The SAA-amplified MSA protofilament structure does closely mimic the structure of one of the protofilaments in the brain-derived MSA α-syn structure, which may explain the similar seeding activities in M83^+/-^ mice ^79^. Other studies have also found differences in the structural or biochemical properties of brain-derived versus *in vitro*-amplified MSA α-syn aggregates ^80,81^. Thus, it is possible that seeding of α-syn aggregation by MSA brain extract in M83^+/-^ mice occurs via a mechanism other than templated conformational conversion. One possibility is that seeding occurs via a secondary nucleation event in which the surface of α-syn fibrils stimulates the formation of new aggregates that do not necessarily exhibit the same structure as the seed ^82–85^. If surface-mediated secondary nucleation dominates during propagation of MSA in M83^+/-^ mice, or if human-specific cellular factors are required to faithfully propagate MSA-associated α-syn aggregates, this may result in a decreased ability of the M83^+/-^ model to discriminate between different α-syn strains. It will therefore be important to test in future studies whether α-syn aggregates from MSA-C and MSA-P also exhibit strain homogeneity in other paradigms.

Currently, there are no approved disease-modifying therapies for MSA, and treatment options are limited to management of symptoms ^86^. Targeting α-syn aggregation and propagation remains an attractive therapeutic strategy for halting disease progression in MSA and other synucleinopathies ^87,88^. As small molecule therapeutics designed to counteract protein misfolding and aggregation can exhibit strain-specific differences in efficacy ^89^, the fact that a single α-syn strain appears to cause both MSA-C and MSA-P may suggest an easier path to developing an α-syn-directed therapeutic that is effective against both MSA clinical subtypes.

## Acknowledgements

The authors are grateful to Ivan Martinez Valbuena and Gabor Kovacs for help with establishing the α-syn seed amplification assay. This work was funded by a grant to J.C.W. from the Canadian Institutes of Health Research (PJT-169042), and J.C.W is supported by a Canada Research Chair (Tier 2) in protein misfolding disorders. H.H.C.L. was supported by the Ernest, Molly and Phillip Giles MSA Research Award. N.R.G.S. was supported by a Parkinson Canada/Parkinson Society of British Columbia doctoral fellowship, a doctoral research award from the Canadian Institutes of Health Research, and an Ontario Graduate Scholarship. S.M. was supported by a postdoctoral fellowship from Parkinson Canada. R.W.L.S. was supported by fellowships from the Croucher Foundation, Parkinson Canada, and the Peterborough K.M. Hunter Charitable Foundation, as well as an Ontario Graduate Scholarship. Experimental schematics were generated using BioRender.com.

## Declarations

### Availability of data and material

All data generated or analyzed during this study are included in this published article or its supplemental information.

### Author contributions

<u>Conceptualization:</u> H.H.C.L, B.T.H., M.I., J.C.W.

<u>Formal analysis and investigation:</u> H.H.C.L, N.R.G.S., S.M., R.W.L.S., L.Y.L, A.M., E.S., C.S.U., J.C.W.

<u>Resources:</u> B.T.H., M.I.

<u>Writing-original draft:</u> H.H.C.L., J.C.W.

<u>Writing-review and editing:</u> H.H.C.L, N.R.G.S., S.M., R.W.L.S., L.Y.L, A.M., E.S., C.S.U., B.T.H., M.I., J.C.W.

<u>Funding acquisition:</u> J.C.W.

<u>Supervision:</u> J.C.W.

## Methods

### Human tissue samples

MSA patient sample details are provided in **Supplemental Table S1**. Frozen tissue was obtained from the temporal cortex, substantia nigra and cerebellum of eight MSA-C and seven MSA-P patients from the Massachusetts Alzheimer’s Disease Research Center. Informed consent was obtained for all cases used in this study at the point of tissue collection. The use of human tissue was conducted in accordance with guidelines provided by the University of Toronto under an approved human participant ethics protocol (#38879; “Use of human tissue for research on neurodegenerative diseases”).

### Generation of human and mouse brain extracts

To prepare the human and mouse brain homogenates used for analysis, frozen tissue was homogenized (10% w/v) in calcium- and magnesium-free PBS using a Minilys homogenizer and CK14 soft tissue homogenizing tubes (Bertin). Homogenates were immediately aliquoted and stored at -80 °C. To generate detergent-extracted brain homogenates, samples were diluted into 1X detergent buffer (0.5% (v/v) Nonidet P-40, 0.5% (w/v) sodium deoxycholate, in PBS) with added Pierce Universal Nuclease (ThermoFisher #88701) and Halt Phosphatase Inhibitor (ThermoFisher #784420). Samples were then incubated on ice for 20 min with occasional vortexing. Mixtures were clarified via centrifugation at 1,000 x g for 5 min at 4 °C and the detergent-extracted supernatants were used for further analysis.

### Immunoblotting

Samples were electrophoresed on 4-12% or 12% Bolt Bis-Tris Plus gels (Thermo Scientific) prior to transfer onto 0.45 μm polyvinylidene fluoride membranes immersed in transfer buffer [25 mM Tris, pH 8.3, 0.192 M glycine, 20% (v/v) methanol] for 1 h at 35 V. Crosslinking was performed by membrane incubation in 0.4% (v/v) paraformaldehyde in PBS for 30 min at room temperature, with rocking ^90^. Membranes were next blocked for 1 h at room temperature with rocking in blocking buffer [5% (w/v) skim milk in 1X TBST (TBS and 0.05% (v/v) Tween-20)] and then incubated overnight at 4 °C with primary antibodies diluted in blocking buffer. The primary antibodies used were anti-α-syn mouse monoclonal Syn-1 (BD Biosciences #610786; 1:10,000 dilution) and anti-PSyn α-syn rabbit monoclonal EP1536Y (Abcam #ab51253; 1:4,000 dilution). Following primary antibody incubation, membranes were washed 3 times with TBST and then incubated, for 1 h at room temperature, with horseradish peroxidase-conjugated secondary antibodies (Bio-Rad #172-1019 or 172-1011) diluted 1:10,000 in blocking buffer. Following another 3 washes with TBST, immunoblots were developed using Western Lightning enhanced chemiluminescence (ECL) Pro (PerkinElmer) or SuperSignal West Dura (ThermoFisher) and imaged using either X-ray film or the LiCor Odyssey Fc system.

### Protease digestion assays

Protein concentration in detergent-extracted brain homogenates was determined using the BCA assay (Thermo Fisher #23225) and then samples were diluted to a concentration of 5 mg/mL total protein using 1X detergent buffer. Samples were digested with either 50 μg/mL thermolysin or 100 μg/mL proteinase K for a total protease/protein (w/w) ratio of 1:100 and 1:50, respectively. Digestions were performed with continuous shaking at 600 rpm at 37 °C on an Eppendorf Thermomixer F1.5 for 1 h and then were stopped by adding EDTA to a final concentration of 5 mM (for TL digestions) or PMSF to a final concentration of 4 mM (for PK digestions). The insoluble fraction was collected by ultracentrifugation in a TLA-55 rotor (Beckman Coulter) at 100,000x g at 4 °C for 1 h, and pellets were resuspended in loading buffer (1X Bolt LDS sample buffer (ThermoFisher Scientific, #B0007) containing 2.5% β-mercaptoethanol) and heated to 95 °C for 10 min. Protease-resistant species were assessed by immunoblotting.

### Conformational stability assays

α-Syn conformational stability assays were performed as previously described ^34,47^. Briefly, GdnHCl was added to an equal amount of detergent-extracted brain homogenate to yield final GdnHCl concentrations of 0, 1, 1.5, 2, 2.5, 3, 3.5 and 4 M in a total volume of 40 µL. Samples were incubated at room temperature with shaking for 2 h (800 rpm) before being diluted to 0.4 M GdnHCl in PBS containing 0.5% (w/v) sodium deoxycholate and 0.5% (v/v) NP-40. Samples were ultracentrifuged at 100,000 x *g* for 1 h at 4 °C in a TLA-55 rotor. Pellets were then resuspended in loading buffer, boiled for 10 min at 95 °C, and then analyzed by immunoblotting. Denaturation curves were generated by performing densitometry on immunoblot images and then normalizing to the highest signal for each sample. The data for independent samples was then averaged and re-normalized so that the value for 0 M GdnHCl was set at 100%. [GdnHCl]_50_ values were calculated by fitting the denaturation curves to a variable slope (four parameter) logarithmic model in GraphPad Prism with the top value fixed at 100 at the bottom value fixed at 0.

### α-Synuclein seed amplification assays

SAAs were performed as previously described ^54^. Briefly, full-length untagged wild-type recombinant human α-syn was purified from *E. coli* using an osmotic shock protocol followed by anion exchange chromatography. Prior to SAA, recombinant α-syn was dialyzed into dH_2_O. PBS-soluble seeds for the SAA experiments were generated by centrifuging 10% (w/v) brain homogenate (in PBS) at 10,000x g for 10 min at 4 °C. The protein concentration in the PBS-soluble fraction was determined using the BCA assay. SAA reactions were performed in black 96-well plates with a clear bottom (Corning). Each well contained 40 mM phosphate buffer pH 8, 175 mM sodium citrate tribasic dihydrate, 10 μM ThT, 0.1 mg/mL recombinant α-syn, 5 μg of PBS-soluble total protein, and three 0.5 mm silica beads in a final volume of 100 µL. Plates were sealed and incubated at 42 °C in a CLARIOstar microplate reader (BMG Labtech) with cycles of 1 min shaking (400 rpm double orbital) and 1 min rest for a total of 72 h. ThT fluorescence measurements (450 ± 10 nm excitation and 480 ± 10 nm emission, bottom read) were taken every 2 min. For each sample, 3-4 technical replicates were performed.

Lag phases for the SAA reactions were calculated by fitting the kinetic curves to a variable slope (four parameter) logarithmic model in GraphPad Prism and then using the equation T_50_ − [1/(2*k)], where k is the Hill slope and T_50_ is the time at which fluorescence is halfway between the baseline and plateau values ^91^. Maximum ThT fluorescence values were calculated as the maximum ThT reading taken during the 72 h assay whereas plateau ThT fluorescence values were calculated as the mean fluorescence during the last hour of the assay.

### Propagation studies in M83^+/-^ mice

To generate the hemizygous M83 mice (M83^+/-^) used for propagation experiments, homozygous M83 transgenic mice on a mixed C57BL6/C3H background (Jackson Lab, #004479) were crossed with non-transgenic B6C3F1 mice (Jackson Lab, #100010). Mice were maintained on a 12 h light and 12 h dark cycle with unlimited access to both food and water and housed in groups of 4-5 animals per cage. Roughly equivalent numbers of female and male mice were used in all inoculation studies. All animal experiments were performed according to guidelines established by the Canadian Council on Animal Care under a University Health Network Animal Care Committee-approved protocol (AUP #4263.23).

Intracerebral inoculations were performed on weanling M83^+/-^ mice at ∼5 weeks of age. Groups of 8-10 mice were anaesthetized using isoflurane gas for sedation prior to inoculation. Non-stereotactic intracerebral inoculations were performed using a tuberculin syringe with an attached 27-gauge, 0.5-inch needle (BD Biosciences #305945), inserted to a depth of ∼3 mm into the right cerebral hemisphere of the brain. The brain regions targeted using this method are the hippocampus and overlying parietal cortex. For inoculations with MSA patient samples, each mouse received 30 µL of 10% (w/v) brain homogenate (in PBS). For the second passage experiments, each mouse received 30 µL of 10% (w/v) brain homogenate (in PBS) from a clinically ill M83^+/-^ mouse that had been previously inoculated with MSA brain extract, or an asymptomatic M83^+/-^ mouse that was previously inoculated with PBS ^34^. Inoculated mice were monitored daily for signs of motor dysfunction (hindlimb paralysis, loss of grip strength, bradykinesia, and weight loss) and were euthanized when paralysis progressed to the point where mice exhibited a reduced ability to ambulate and obtain food. Euthanization was performed via intraperitoneal injection of sodium pentobarbital followed by transcardiac perfusion with 0.9% saline solution, and then brains were collected and divided parasagitally with the left hemisphere frozen at -80 °C and the right hemisphere fixed in 10% neutral buffered formalin.

### Neuropathological analysis

Immunohistochemical staining for cerebral α-syn pathology was performed as previously described ^35^. Briefly, formalin-fixed hemibrains were embedded in paraffin, sectioned into 5 µm sagittal slices, and then deparaffinized and rehydrated. Epitope retrieval was performed in citrate buffer [10 mM sodium citrate, pH 6, with 0.05% (v/v) Tween-20] for 15 min in a pressure cooker (Instant Pot, #112-0141-01). Endogenous peroxidase activity was inhibited by incubation in 3% H_2_O_2_ for 5 min, and then slides were blocked with 2.5% (v/v) normal horse serum (Vector Laboratories, #MP-7401) at 22 °C for 1 h. Slides were stained with the anti-PSyn antibody EP1536Y (Abcam, #ab51253) diluted 1:320,000 in DAKO antibody diluent (Agilent, #S0809) at 4 °C overnight. For the serial passaging experiments, an antibody dilution of 1:10,000 was used. After washing, slides were incubated in secondary antibody [ImmPress horseradish-peroxidase-labeled horse anti-rabbit detection kit (Vector Laboratories, #MP-7401)] at 22 °C for 30 min and then developed using ImmPACT 3,3’-diaminobenzidine peroxidase substrate (Vector Laboratories, #SK-4105). Slides were then counterstained with hematoxylin (Sigma-Aldrich, #GHS132) and mounted using Cytoseal 60 (Epredia, #8310–4) then scanned with the TissueSnap and TissueScope LE120 systems (Huron Digital Pathology) and visualized with PMA.start (Pathomation). The area covered by PSyn staining in the midbrain, hypothalamus, and medulla of MSA-inoculated M83^+/-^ mice was quantified using ImageJ as previously described using a threshold range of 0-130 ^20,35,48^. The number of PSyn-positive cells in the cerebral cortex was counted manually.

### Statistical analysis

All statistical analysis was performed using GraphPad Prism (version 10.6.1). The threshold for statistical significance was set at P < 0.05. Data was not assumed to be normally distributed.

Survival curves following inoculation of M83^+/-^ mice were compared using the Log-rank test. The extent of PSyn deposition in different brain regions of inoculated M83^+/-^ mice was compared by two-way ANOVA followed by Šidák’s multiple comparisons test. The Mann-Whitney test was used for all other statistical comparisons.

---

**Supplemental Table S1.**
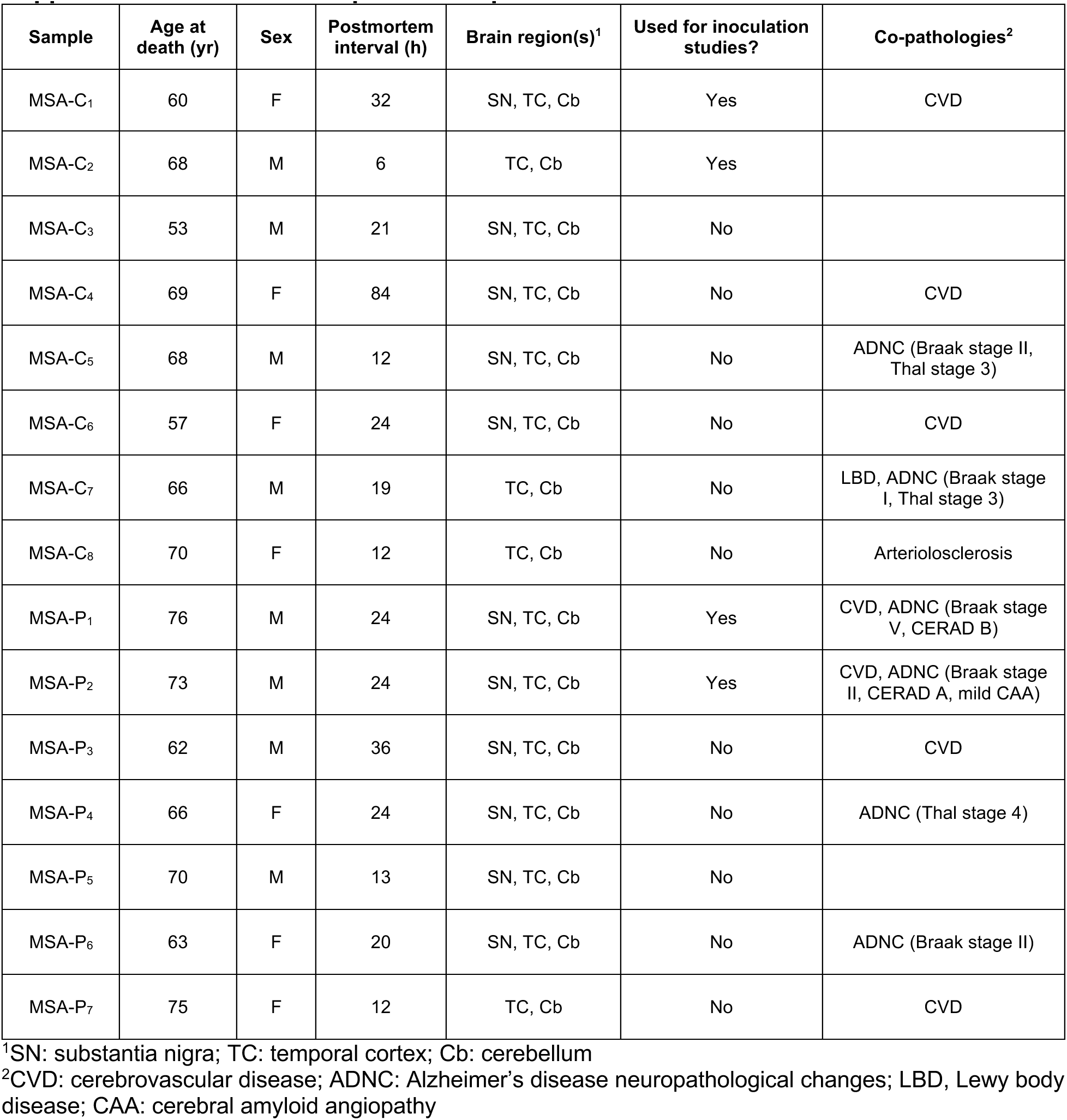
MSA patient sample details.

**Supplemental Figure S1.**
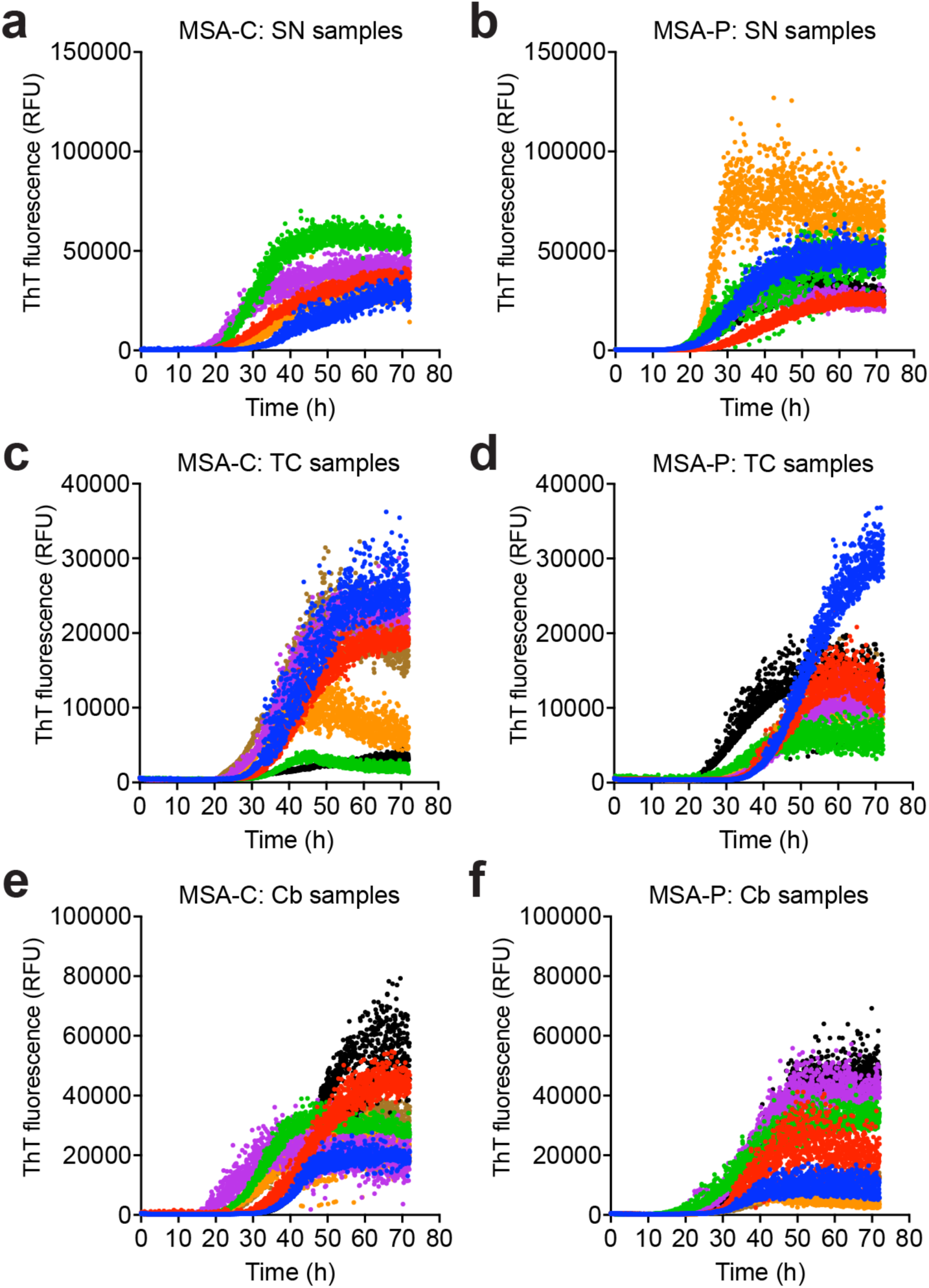
α-Synuclein seed amplification curves for individual MSA-C and MSA-P patient samples. Individual ThT aggregation curves for SAAs using homogenates from the substantia nigra (a, b), temporal cortex (c, d), or cerebellum (e, f) of MSA-C (a, c, e) and MSA-P (b, d, f) patients. For SN samples, n = 5 (MSA-C) or n = 6 (MSA-P); for TC and Cb samples, n = 7 per MSA subtype. Each curve depicts the mean ThT fluorescence over time for 3-4 technical replicates per MSA sample.

**Supplemental Figure S2.**
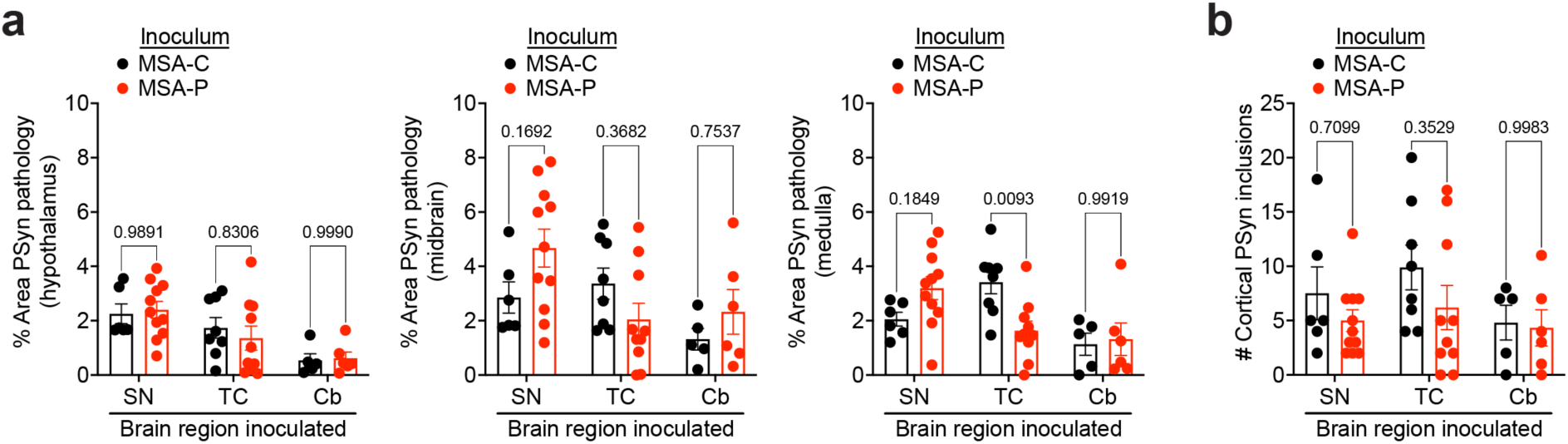
Quantification of α-syn deposition in the brains of M83^+/-^ mice inoculated with samples from different brain regions of MSA-C and MSA-P patients. **a**) Quantification of the percent area covered by PSyn staining in the hypothalamus (left graph), midbrain (middle graph), or medulla (right graph) from M83^+/-^ mice inoculated with substantia nigra, temporal cortex, or cerebellar homogenate from MSA-C or MSA-P patients. **b**) Quantification of the number of cortical PSyn deposits in the brains of M83^+/-^ mice inoculated with substantia nigra, temporal cortex, or cerebellar homogenate from MSA-C or MSA-P patients. For SN samples, n = 6 or 11 mice for MSA-C and MSA-P, respectively; for TC samples, n = 8 or 10 mice for MSA-C and MSA-P, respectively; for Cb samples, n = 5 or 6 mice for MSA-C and MSA-P, respectively. Graphs display mean ± s.e.m. Statistical significance was assessed by two-way ANOVA followed by Šidák’s multiple comparisons test.

**Supplemental Figure S3.**
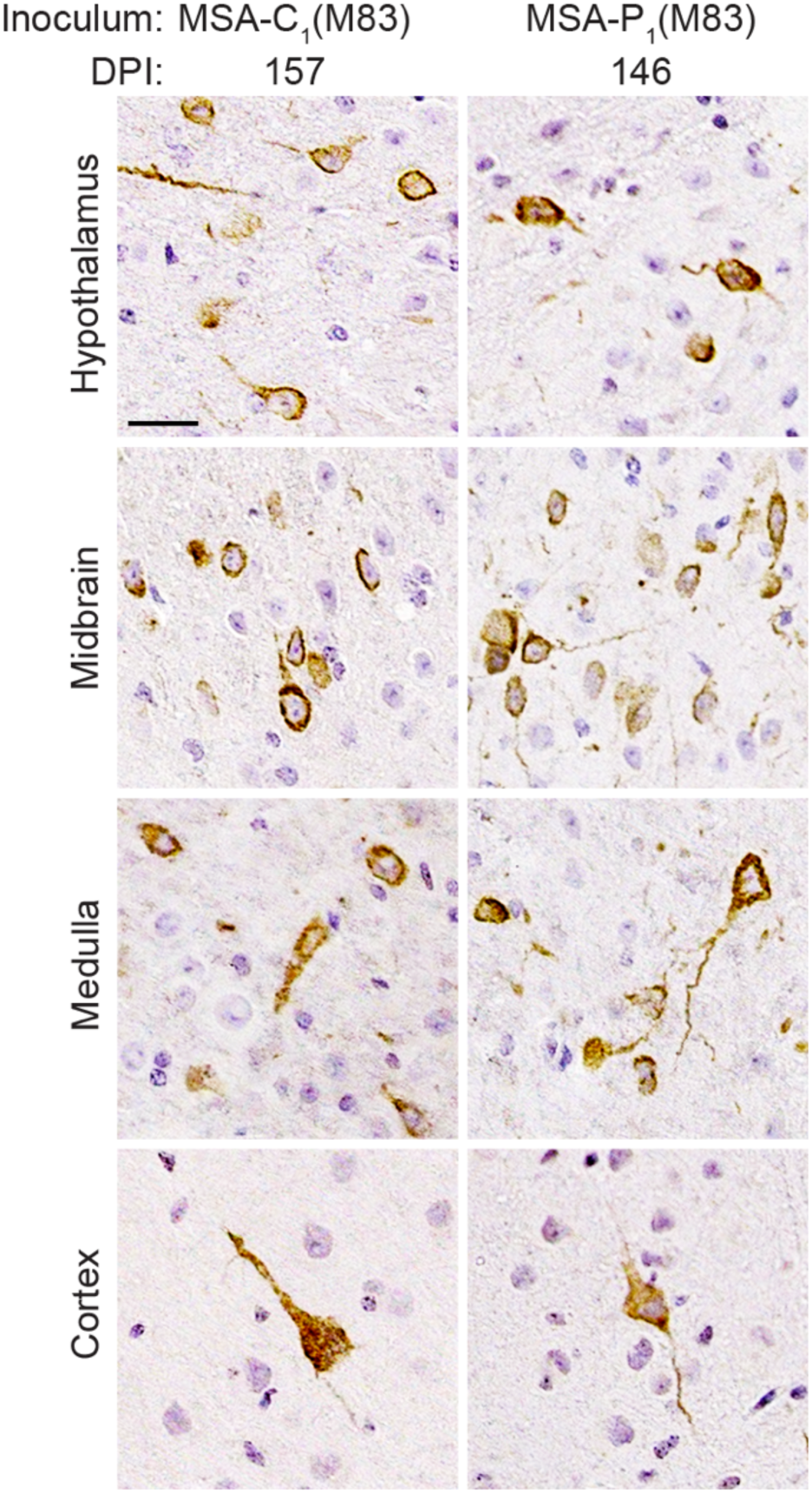
Cerebral α-syn deposition in M83^+/-^ mice inoculated with passaged MSA-C and MSA-P samples. Representative images of PSyn-stained sections from the hypothalamus, midbrain, medulla, and cortex of clinically ill M83^+/-^ mice following inoculation with the indicated passaged MSA samples. PSyn was detected using the antibody EP1536Y. Scale bar = 20 µm (applies to all sections).

